# Optimizing the delivery of self-disseminating vaccines in fluctuating wildlife populations

**DOI:** 10.1101/2022.12.13.520205

**Authors:** Courtney L. Schreiner, Andrew J. Basinski, Christopher H. Remien, Scott L. Nuismer

**Affiliations:** Department of Ecology and Evolutionary Biology, University of Tennessee Knoxville; Institute for Interdisciplinary Data Sciences, University of Idaho; Department of Mathematics and Statistical Science, University of Idaho; Department of Biological Sciences, University of Idaho

## Abstract

Zoonotic pathogens spread by wildlife continue to spill into human populations and threaten human lives. A potential way to reduce this threat is by vaccinating wildlife species that harbor pathogens that are infectious to humans. Unfortunately, even in cases where vaccines can be distributed en masse as edible baits, achieving levels of vaccine coverage sufficient for pathogen elimination is rare. Developing vaccines that self-disseminate may help solve this problem by magnifying the impact of limited direct vaccination. Although models exist that quantify how well these self-disseminating vaccines will work when introduced into temporally stable wildlife populations, how well they will perform when introduced into populations with pronounced seasonal population dynamics remains unknown. Here we develop and analyze mathematical models of fluctuating wildlife populations that allow us to study how reservoir ecology, vaccine design, and vaccine delivery interact to influence vaccine coverage and opportunities for pathogen elimination. Our results demonstrate that the timing of vaccine delivery can make or break the success of vaccination programs. As a general rule, the effectiveness of self-disseminating vaccines is optimized by introducing after the peak of seasonal reproduction when the number of susceptible animals is near its maximum.

## 2 Introduction

The majority of human infectious diseases are caused by pathogens with animal origins (Jones et al., 2008). As the human population continues to encroach on wildlife habitat, zoonotic pathogens such as Ebola virus, *Borrelia burgdorferi*, Lassa virus, Sin Nombre virus, and Nipah virus pose an increasing threat of spillover into the human population (Gottdenker et al., 2014; Pongsiri et al., 2009; Keesing et al., 2010; Coltart et al., 2017; Jones et al., 2008). Several of these emerging infectious diseases have had devastating impacts on public health. The 2014 Ebola outbreak, for example, killed more than 11,000 people (Coltart et al., 2017), and the ongoing SARS-CoV-2 pandemic has killed millions (WHO, 2021). The SARS-CoV-2 pandemic has made the perils of our current reactionary approach to managing emerging infectious disease clear and helped to focus attention on methods that proactively reduce the risk of spillover and emergence.

Vaccinating wildlife reservoir populations is a proven method for lowering pathogen prevalence and reducing the risk of spillover into the human population (Hampson et al., 2007; Velasco-Villa et al., 2017). For example, oral rabies vaccines that are distributed in bait-form have proven to be effective at controlling rabies in fox and raccoon populations (Freuling et al., 2013; Sidwa et al., 2005; MacInnes et al., 2001). However, even in these cases where an effective bait-deliverable vaccine exists, it remains difficult to achieve a level of vaccination coverage sufficient for pathogen elimination (Ramey et al., 2008; Sattler et al., 2009). The key obstacles are the cost and logistical difficulty of distributing vaccine into inaccessible wildlife populations. For zoonotic infectious diseases with short-lived reservoirs (e.g., rodents), the challenge is compounded by the rapid dilution of immunity established through traditional vaccination. These challenges suggest that distributing traditional vaccines as baits is unlikely to provide a general solution (Nuismer et al., 2020; Mariën et al., 2019).

Recent developments in vaccine design offer fresh solutions to this long-standing problem by creating vaccines that are capable of some degree of self-dissemination. Self-disseminating vaccines can be either transferable or transmissible. Development of transferable vaccines has focused on applying topical vaccine-laced gels to individual animals (Bakker et al., 2019). When other individuals engage in natural allogrooming behaviors common in some reservoir species (e.g., bats), they ingest the vaccine and gain immunity. As a result, the number of animals that can be vaccinated is substantially multiplied (Bakker et al., 2019). In contrast to transferable vaccines which do not generate sustained chains of self-dissemination, transmissible vaccines are engineered to be contagious, and are potentially capable of indefinite self-dissemination within the reservoir population (Nuismer and Bull, 2020). A diverse range of modeling studies have demonstrated that both types of self-disseminating vaccines reduce the effort required to achieve herd immunity within wildlife reservoir populations (Nuismer and Bull, 2020; Bakker et al., 2019; Nuismer et al., 2016; Layman et al., 2021; Varrelman et al., 2019; Basinski et al., 2018, 2019). We do not yet know, however, how the introduction of these vaccines can be best timed to maximize their impact when used in reservoir species that have pronounced seasonal population dynamics.

Previous modeling work has demonstrated that the success of traditional wildlife vaccination campaigns can be improved by timing vaccine introduction to coincide with seasonal birth pulses in short-lived animal species (Schreiner et al., 2020). Although intuition suggests similar results should hold for self-disseminating vaccines, the quantitative details remain unknown and important questions remain unanswered. For instance, is timing vaccine introduction more important in transferable vaccines than transmissible vaccines? Do the detailed transmission dynamics of the vaccine (e.g., transmission rate and duration of self-dissemination) influence the optimal timing of introduction? Does timing matter more for some reservoir species than others? Here we develop a general mathematical modeling framework for transmissible and transferable vaccines and use it to quantify the consequences of introducing self-disseminating vaccines at different times throughout the year. We then apply our model to two specific reservoir species that harbor important human pathogens: the primary reservoir of Lassa virus, *Mastomys natalensis*, more commonly known as the multimammete rat and an important carrier of Rabies virus, *Desmodus rotundus*, frequently referred to as the common vampire bat. The specific questions we address are: 1) What is the optimal time of year to distribute a self-disseminating vaccine? 2) In which situations is optimal timing critical for success? 3) How does the duration of self-dissemination affect the optimal vaccination strategy? 4) How does host demography influence the importance of timing vaccine distribution?

## 3 Methods

We use an SIR (Susceptible-Infected-Recovered) modeling framework to study how the timing of vaccination influences the ability of a self-disseminating vaccine to protect a population from a pathogen. We focus our efforts on populations that undergo seasonal fluctuations in population density driven by well-defined seasonal patterns of reproduction. Our models assume vaccines are introduced into relatively small geographic areas within which the reservoir population is well mixed and of modest size (e.g., 2000 individuals). These assumptions are motivated by rodent species such as *Mastomys natalensis* and *Peromyscus maniculatis* that harbor important human pathogens such as Lassa virus and Sin Nombre virus, respectively (Leirs et al., 1994; Luis et al., 2010).

In the model, we use a time-dependent birth function that is a variation of the periodic Gaussian function developed by Peel et al. (2014):

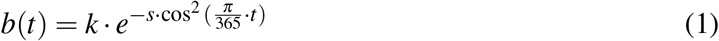

where *s* tunes the synchrony of births, *k* is set so that the average annual population size is equal to 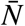, and time is measured in units of days (see Appendix for more details).

Direct vaccination is assumed to occur each year beginning *t_v_* days after the start of the reproductive season and continue for *V_l_* days. Assuming *N_v_* vaccine-laced baits are distributed each year (transmissible vaccine) or *N_v_* animals are painted with vaccine-laced gel (transferable vaccine) at a rate *σ*(*t*), the rate at which individuals are directly vaccinated is given by:

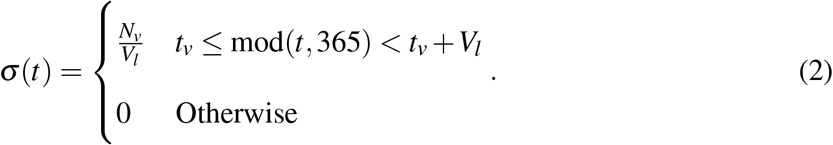

### 3.1 Transmissible vaccine model

Our transmissible vaccine model contains four classes: individuals that are susceptible to both the pathogen and the vaccine (*S*), individuals that are infected with the pathogen (*P*), vaccinated individuals that are immune to the pathogen and capable of transmitting vaccine to susceptible individuals (*V*), and individuals that have immunity due to recovery from pathogen infection or from vaccination (*R*). For simplicity, we assume individuals that have recovered from either the pathogen or the vaccine maintain lifelong immunity to both, and that co-infection with vaccine and pathogen does not occur. Individuals that are infected with the pathogen recover at rate *γ_P_*, and individuals infected with the vaccine recover at rate *γ_V_*. We assume density-dependent transmission of the pathogen and the vaccine, with transmission coefficients *β_P_* and *β_V_* respectively. Individuals may also be lost from the system due to pathogen-induced mortality at rate *v*. Setting the transmission rate of the vaccine *β_V_* equal to zero yields a model for a traditional vaccination campaign.

Susceptible individuals can be vaccinated directly or by coming into contact with vaccine-infected individuals. Because vaccine-laced baits can be consumed by any individual in the population, including individuals already immune to the pathogen, waste is inevitable. We model this feature of vaccine distribution by multiplying the rate at which vaccines are deployed at time t, *σ*(*t*), by the fraction of susceptible individuals 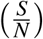 in the population. Here, *N* denotes the total population size. Thus, if the entire population is susceptible, vaccination efficiency is high and waste is low. In contrast, if the population contains a large proportion of immune individuals, vaccination efficiency is low and waste is high. A description of all parameters can be found in Table 1. Together, these assumptions lead to the following system of differential equations:

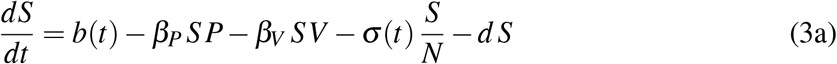

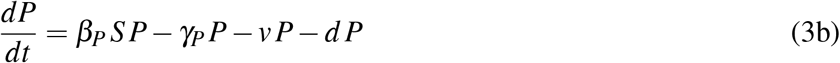

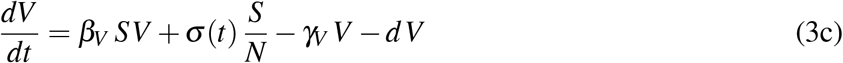

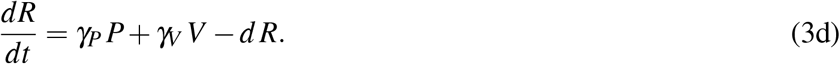

### 3.2 Transferable vaccine model

Our transferable vaccine model contains five classes: individuals that are susceptible to the pathogen (*S*), individuals that are currently infected by the pathogen (*P*), individuals that are immune to the pathogen (*R*), individuals that are currently infected by the pathogen and also carrying the vaccine-laced topical gel (*P_g_*), and individuals that are immune to the pathogen and also carrying the vaccine laced topical gel (*R_g_*). We assume vaccine-laced gel is applied topically to captured animals at rate *σ* (*t*). These animals are also assumed to be directly vaccinated upon capture so that susceptible individuals immediately transition to the *R_g_* class. In contrast to the transmissible vaccine model, the rate of vaccination is multiplied by 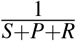 rather than 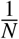. This is because we assume that if individuals have gel on them, it will be recognized and additional gel will not be applied and wasted. Allogrooming behavior allows an individual to become vaccinated at rate *β_g_* if it encounters an individual carrying the vaccine-laced gel. At the same time, however, allogrooming behavior also depletes the quantity of vaccine-laced gel on individual carriers. We model this phenomenon by assuming the topical gel is lost at rate *αN* which implies gel is lost more rapidly in densely populated animal populations. Additionally, we assume the topical gel loses its ability to serve as a vaccine over time at rate *γ_g_*.

We assume that transfer of the vaccine can occur only from an individual to which vaccine-laced gel has been directly applied and that vaccine transfer is density dependent. Pathogen transmission is also assumed to be density-dependent and to occur at rate *β_P_* from contact with either a pathogen-infected individual (*P*) or a gelled and pathogen-infected individual (*P_g_*). See Table 1 for parameter descriptions. Together, these assumptions lead to the following system of differential equations:

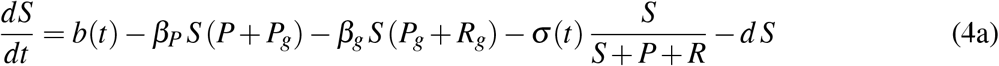

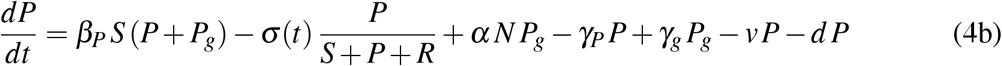

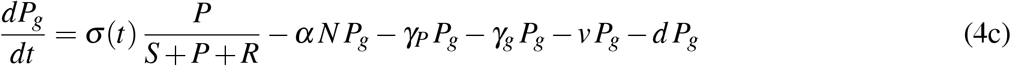

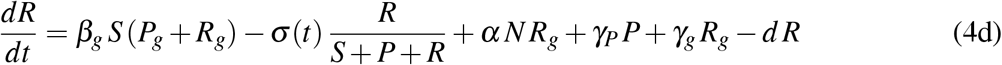

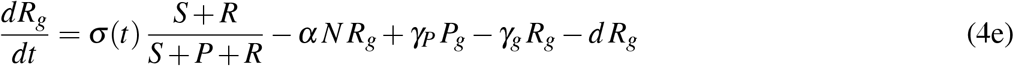

### 3.3 Assessment of vaccination strategy

We evaluate the success of a vaccination campaign by comparing the reduction of pathogen-infected individuals it achieves relative to the situation where no vaccination occurs. For each type of vaccine and distribution strategy, we use the deSolve package in R to numerically solve the corresponding system of differential equations (Soetaert et al., 2010). For each combination of parameters we solve the system of differential equations twice: once with vaccination and once without vaccination. Initial conditions are identical for these two cases and both are burned in for 100 years, allowing the system to settle into stable seasonal cycles. One numerical solution is continued from this point for ten years with no vaccination occurring and the other is run with vaccination for ten years after the first day of vaccination. We then extract from each of the numerical solutions the average number of pathogen-infected hosts over the ten year period following the burn-in. Specifically, we calculate the fractional reduction of pathogen-infected individuals (average level of pathogen reduction) provided by vaccination as:

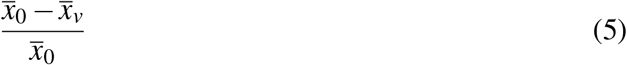

where 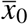 is the average number of pathogen-infected individuals in the scenario without vaccination and 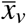 is the average number of pathogen-infected individuals with vaccination. We use this comparative approach to explore how the benefits of vaccination change as a function of vaccine properties, reservoir properties, and the timing of vaccine introduction. Additionally, we use the concept of the basic reproductive number, denoted as *R*_0_, to compare the relative transmissibility of the vaccine and the pathogen. *R*_0_ represents the average number of new infections caused by a single infected individual that is introduced into a fully susceptible population (Keeling and Rohani, 2011). More details on the *R*_0_ calculations for transmissible and transferable vaccines can be found in the Appendix.

### 3.4 Case studies

Up to this point we have developed general models to explore a wide range of parameter space. Our goal was to develop a general understanding of the performance of self-disseminating vaccines as a function of reservoir biology, vaccine properties, and introduction protocol. Next, we shift our focus to specific hosts and the pathogens they carry. We use estimates from the literature to parameterize our model and draw conclusions for two specific systems where self-disseminating vaccines are being developed. Specifically, we focus on the primary rodent reservoir of Lassa virus, *Mastomys natalensis* and a bat reservoir of Rabies virus *Desmodus rotundus*. A list of parameters used in both the general simulations and specific case studies can be found in the supplemental material.

#### 3.4.1 *Mastomys natlensis* – Lassa virus

Our first case-study is the primary rodent reservoir of Lassa virus, *M. natalensis*. Lassa virus commonly spills over into the human population through rodent droppings and leads to the development of Lassa fever which can be fatal in humans (McCormick et al., 1987; Dan-Nwafor et al., 2019). Population sizes of *M. natalensis*, have been shown to fluctuate seasonally in response to birth pulses coinciding with the beginning of the wet season and an increase in the availability of green grass as well as other food sources (Leirs et al., 1997; McCormick et al., 1987). We use data from a study in Guinea – where Lassa virus is endemic – to estimate the level of seasonality that these populations demonstrate (Fichet-Calvet et al., 2007). We use a population size of 2000 as estimated by Mariën et al. (2019). Additionally, parameters estimated from Nuismer et al. (2020) suggest a lifespan of one year for the rodent reservoir, a rate of recovery from Lassa virus infection equal to 21 days, and a Lassa virus *R*_0,*P*_ = 1.5. We are then able to solve for the transmission coefficient *β_P_* based on *γ_p_* and *R*_0,*P*_ (Appendix). We base the transmissible vaccine parameters on a recent study (Varrelman et al., 2022) which suggests that the rodents would be infectious with the vaccine for their entire life (*γ_v_* = 0). We consider a range of values for the reproductive number of the vaccine (*R*_0,*V*_) and we use this predefined *R*_0,*V*_ as well as the recovery rate to calculate the transmission rate of the vaccine (Appendix).

#### 3.4.2 *Desmodus rotundus* – Rabies virus

Our second case-study focuses on the vampire bat, *D. rotundus*, which serves as a reservoir for rabies virus within Central and South America. Rabies is a disease caused by *Rabies lyssavirus* commonly spread by bats and is fatal in most mammals, including humans (Fisher et al., 2018). Vampire bats show evidence of seasonal births and previous studies have used lactation rates to estimate the reproductive seasonality in these populations (Blackwood et al., 2013). We tailor our birth function to data on lactation from Lord (1992) (see Appendix). Although local population sizes of *D. rotundus* are unclear, estimates for colony size do exist. For this reason we focus on a vaccination campaign targeting a single colony of 240 individuals as estimated by Bakker et al. (2019). Estimates suggest that *D. rotundus* live for an average of three and a half years (Lord et al., 1976). To simulate the pathogen dynamics of Rabies we use a pathogen *R*_0_ of 1.5 and an average duration of infection of 21 days (Blackwood et al., 2013; Hampson et al., 2009; Moreno and Baer, 1980). Because roughly 10% of bats that are exposed to rabies end up developing a lethal infection (Blackwood et al., 2013; Bakker et al., 2019), we assume individuals infected with the pathogen have a 10% chance of dying due to infection.

Because both transferable and transmissible vaccines are currently being developed for *D. rotundus* we study both scenarios. Specifically we assume the transferable vaccine gel stays on for approximately two days (*γ_g_* = 1/2) as suggested by Bakker et al. (2019). For the transmissible vaccine, because the proposed transmissible vaccine vector is a betaherpesvirus we assume the vaccine will induce lifelong infection (*γ_v_* = 0) (Griffiths et al., 2020). It is unclear what *R*_0_ these self-disseminating vaccines will have, thus, we explore a range of vaccine *R*_0_ values.

**Table 1:** Model parameters and biological interpretation. Parameter values used in simulations can be found in the supporting online material.

| Parameter list |  |
| --- | --- |
| Parameter | Description |
| $t_v$ | Day in year of vaccine initiation |
| $V_l$ | Duration of the vaccination campaign (days) |
| $s$ | Synchrony of births |
| $d$ | Natural mortality rate (per individual per day) |
| $\bar{N}$ | Average population size |
| $R_{0,V}$ | $R_0$ of the vaccine |
| $R_{0,P}$ | $R_0$ of the pathogen |
| Parameter | Description |
| $\gamma_P$ | Recovery rate of the pathogen (per individual per day) |
| $\gamma_V$ | Recovery rate of the transmissible vaccine (per individual per day) |
| $\gamma_g$ | Recovery rate of the transferable vaccine (per individual per day) |
| $\beta_P$ | Rate of pathogen transmission (per individual per day) |
| $\beta_V$ | Rate of transmissible vaccine transmission (per individual per day) |
| $\beta_g$ | Rate of transferable vaccine transmission (per individual per day) |
| $\nu$ | Rate of pathogen induced mortality (per individual per day) |
| $\alpha$ | Rate at which individuals remove gel via grooming (per individual per day) |

## 4 General results

### 4.1 Temporal dynamics of immunity depend on the type of self-disseminating vaccine

Previous work has demonstrated that self-dissemination increases vaccine coverage and reduces the effort required for pathogen elimination (Nuismer and Bull, 2020). However, it remains unclear how self-disseminating vaccines will perform in fluctuating populations. To establish baseline expectations for the performance of self-disseminating vaccines in fluctuating reservoir populations we begin by studying the dynamics of immunity in the absence of the pathogen. Numerical analyses performed over a wide range of parameters demonstrate that the temporal dynamics of immunity differ across vaccine types in characteristic ways (Figure 1). For conventional vaccines that lack the ability to self-disseminate, vaccination results in a rapid increase in the number of vaccinated individuals, followed by a decrease due to the continued influx of susceptible individuals during the birthing season. Transferable vaccines result in similar temporal dynamics but show a transient increase in immunity from self-dissemination following vaccine introduction. In contrast, transmissible vaccines with an *R*_0,*V*_ > 1 can continue to increase the number of immune individuals long after vaccine introduction because they generate self-sustaining chains of transmission. Because all individuals die at a constant rate *d*, the number of immune individuals decreases after the birth pulse ends until the next vaccination campaign for all types of vaccine. With self-disseminating vaccines, the level of increase in the number of immune individuals in the population is dependent on the vaccine *R*_0_ (*R*_0,*V*_) (Figure 1).

**Figure 1:**
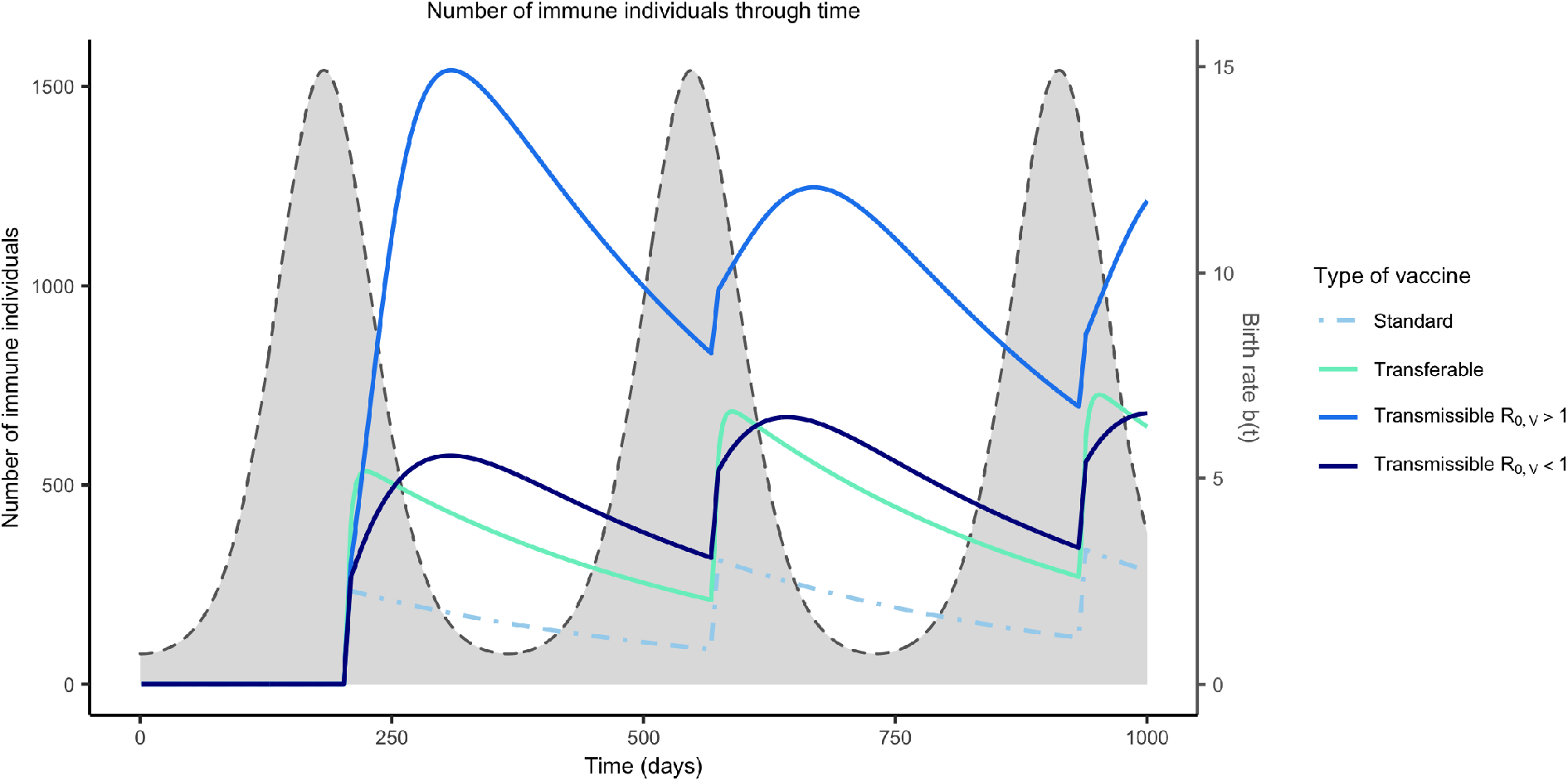
The temporal dynamics of immunity for standard, transferable, and transmissible vaccines in the absence of a pathogen. For each type of vaccine, 250 vaccines are distributed on day 200. The colored lines represent the number of immune individuals in the population over three years of repeated vaccination for either a standard vaccine, transferable vaccine, transmissible vaccine with *R*_0,*V*_ < 1, and a transmissible vaccine with *R*_0,*V*_ > 1. *R*_0,*V*_ of the standard, transferable, strongly transmissible, and weakly transmissible are: (0, 1.5, 1.5, and 0.75) respectively. The remaining parameters are: an average population size of 2000 individuals 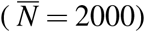, *s* = 3, an average lifespan of 1 year (*d* = 1/365), *R*_0,*P*_ = 2, 250 vaccines are distributed each year (*N_V_* = 250), individuals can disseminate vaccine for 21 days on average (*γ_V_* = 21^−1^), individuals remain infectious with the pathogen for 21 days on average (*γ_P_* = 21^−1^), the transferable vaccine is groomed off individuals after 6 days on average (*α* = 1/15000, and the pathogen is non-virulent (*v* = 0).

### 4.2 Timing is critical for most self-disseminating vaccines

Previous work has shown that the timing of delivery for conventional vaccines matters in short-lived animals with distinct reproductive seasons (Schreiner et al., 2020). Here, our goal is to evaluate whether timing is more important for transmissible or transferable vaccines and under which conditions timing matters most. To this end, we compared the reduction in pathogen prevalence achieved for vaccination campaigns that are initiated at different times of year and last for various lengths of time. Our results demonstrate that distributing self-disseminating vaccines slightly after the peak of the birthing season will substantially reduce pathogen prevalence (Figure 2). This occurs because it is at this time that population density and the proportion of susceptible individuals are near their seasonal maxima. This ensures that vaccines are not wasted by distributing vaccine at the wrong time. If, however, a large number of vaccines are available and can be distributed, a greater level of pathogen reduction can be achieved and the importance of timing decreases (Supplemental Figure 1).

**Figure 2:**
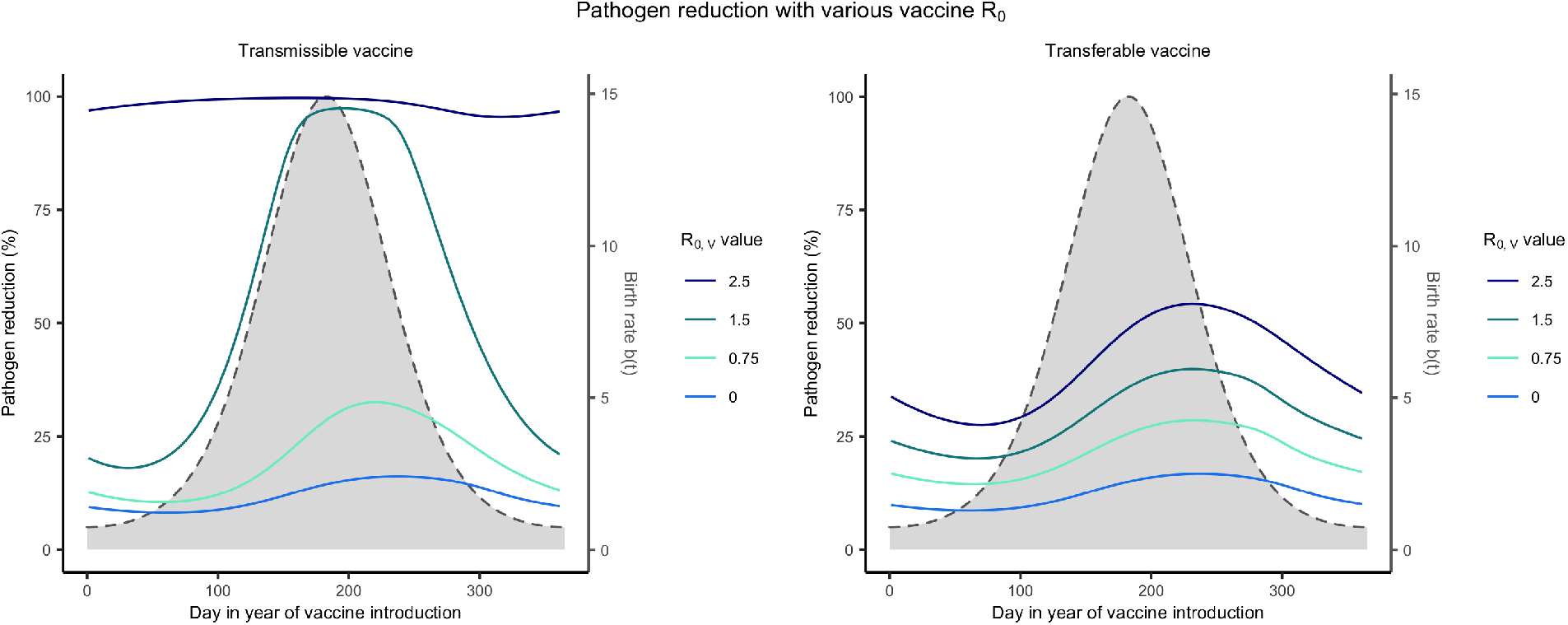
Optimal timing for self-disseminating vaccines as a function of vaccine *R*_0,*V*_. Solid lines represent the level of pathogen reduction achieved for a given date of vaccine introduction for different vaccine *R*_0,*V*_. The grey region outlined by the dashed lined represents the seasonal birthing season where day 1 corresponds to the first day of the birthing season. Additional parameters used were: an average population size of 2000 individuals 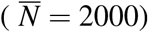, *s* = 3, an average lifespan of 1 year (*d* = 1/365), *R*_0,*P*_ = 2, 250 vaccines are distributed each year (*N_V_* = 250), individuals can disseminate vaccine for 21 days on average (*γ_V_* and *γ_g_* = 21^−1^), individuals remain infectious with the pathogen for 21 days on average (*γ_P_* = 21^−1^), the transferable vaccine is groomed off individuals after 6 days on average (*α* = 1 /15000, and the pathogen is non-virulent (*v* = 0).

For both types of self-disseminating vaccine, pathogen reduction is greater with a larger vaccine *R*_0_. In addition to facilitating pathogen elimination, increasing the transmissible vaccine’s *R*_0,*V*_ also increases the range of times over which a vaccine can be introduced and still substantially reduce the pathogen’s prevalence (Figure 2). This occurs because increased transmission allows the vaccine to be introduced earlier in the reproductive season and still reach individuals that will be born later through downstream transmission. In contrast, with reduced transmission (lower *R*_0,*V*_), if a transmissible vaccine is introduced too early, chains of transmission are generally too short to reach individuals born later in the season resulting in wasted vaccine. Once the *R*_0,*V*_ of the transmissible vaccine exceeds that of the pathogen *R*_0,*P*_, timing matters little and significant pathogen reductions can be accomplished for a broad range of introduction times (Figure 2). This is because a vaccine more transmissible than the target pathogen can out-compete the pathogen and will inevitably displace it from the population over time (Nuismer et al., 2016). A fundamental difference for transferable vaccines is that they never reach this same level of insensitivity to the timing of introduction. The reason for this is that they are (by definition) capable of spreading only from individuals that have been directly vaccinated and thus generate chains of transmission only one step long. Because of this limited spread, an increased *R*_0,*V*_ of the transferable vaccine results in higher levels of pathogen reduction, but not an increase in the range of times over which high pathogen reduction can be achieved (Figure 2).

In general, self-disseminating vaccines should be distributed after the peak of the birthing season to maximize their impact. Specifically, transferable vaccines cause the greatest reduction in the number of pathogen-infected individuals when introduced after the peak of the birthing season. In contrast, transmissible vaccines cause the greatest reduction in the number of pathogen-infected individuals when introduced during the birth pulse, with the optimal solution depending on vaccine *R*_0_. Specifically, the impact of transmissible vaccines with intermediate *R*_0,*V*_ is maximized by early introduction. This occurs because these highly transmissible vaccines can be introduced when newly born susceptible individuals are relatively rare and yet still reach susceptible individuals born later. In contrast, transmissible vaccines with small *R*_0,*V*_ must be introduced later and after a significant number of susceptible individuals has accumulated in order to persist and spread (Figure 2).

For vaccination campaigns of feasible duration (one week - 2 months), the duration of the vaccination campaign itself matters little as long as the total amount of distributed vaccine is held fixed (Figure 3). This insensitivity arises primarily because birth rates change little over such short periods of time in most systems. In special cases where it is possible to distribute vaccine over greater periods of time, differences do begin to develop (Figure 3 vertical axis). Generally a longer vaccination campaign results in a lower overall vaccination rate because vaccines are distributed when few susceptible individuals exist within the reservoir population and are thus wasted. If, however, the vaccination campaign begins at the wrong time (i.e., after the birthing season), extending the duration of vaccine-delivery can compensate to some degree (Figure 3). If the timing of birthing within the reservoir population is known, however, the best solution for maximizing the reduction in pathogen prevalence is to distribute vaccines shortly after the peak of the birthing season and over a relatively short amount of time.

**Figure 3:**
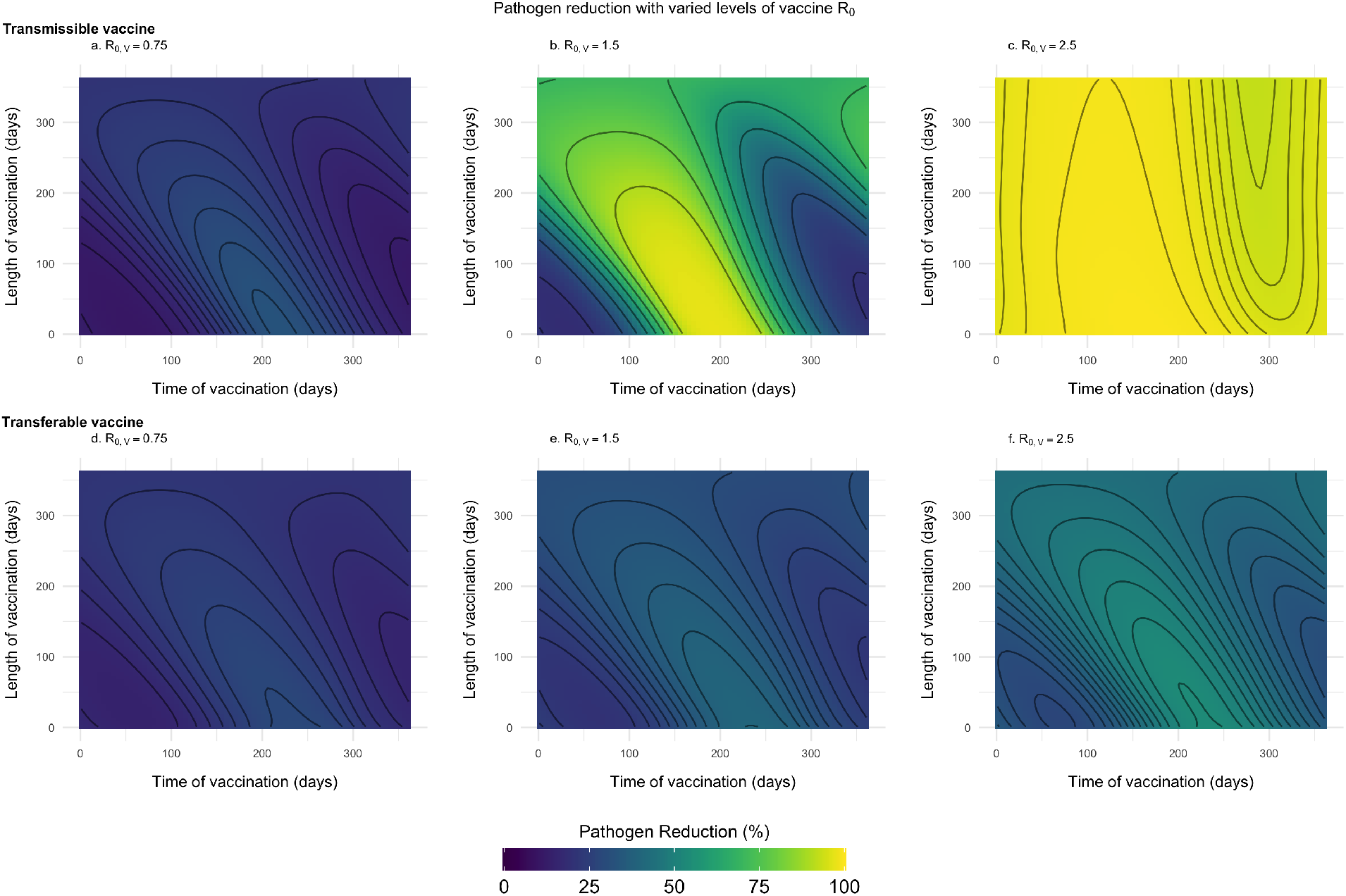
Level of pathogen reduction achieved for both transmissible vaccines and transferable vaccines at different times and for different durations of a vaccination campaign. The R0 in the figure refers to the vaccine *R*_0_. The remaining parameters used were: an average population size of 2000 individuals 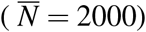, *s* = 3, an average lifespan of 1 year (*d* = 1/365), *R*_0,*P*_ = 2, 250 vaccines are distributed each year (*N_V_* = 250), individuals can disseminate vaccine for 21 days on average (*γ_V_* and *γ_g_* = 21^−1^), individuals remain infectious with the pathogen for 21 days on average (*γ_P_* = 21^−1^), the transferable vaccine is groomed off individuals after 6 days on average (*α* = 1/15000, and the pathogen is non-virulent (*v* = 0).

### 4.3 Vaccines with temporally focused self-dissemination are more effective

Because vaccines may differ widely in the period of time over which they self-disseminate, we explored how this property influenced the optimal timing of delivery. For both types of vaccines, we considered scenarios where the vaccine self-disseminated for 14, 21, 30, 182, and 365 days on average, with vaccine *R*_0_ held constant at a value of 1.5. Holding *R*_0,*V*_ constant while changing the duration of self-dissemination requires that the rate of vaccine transmission also changes *β_V_*. Thus, vaccines with temporally focused periods of self-dissemination also have a high transmission rate whereas vaccines with drawn out periods of self-dissemination have a low transmission rate. If, however, the vaccine *R*_0_ is not held constant by changing the rate of vaccine transmission, then increasing the duration of self dissemination increases vaccine *R*_0,*V*_ leading to higher levels of pathogen reduction.

Our results indicate that vaccines that disseminate for short periods of time are more effective and create greater opportunity for pathogen reduction (Figure 4). Transferable vaccines achieve the highest level of pathogen reduction with acute durations of self-dissemination. This is because with long durations of self-dissemination, *β_V_* is smaller and thus it takes longer to infect individuals with the vaccine. These slow dynamics of the vaccine cause transferable vaccines to miss the peak of the birthing season. However, since the transferable vaccine is groomed off of individuals at rate (*α*), the lengths of self-dissemination that are longer than the average duration gel remains on individuals show no difference (Figure 4). The reverse is also found if we compare different alpha values (Supplemental Figure 2). In contrast, the transmissible vaccine can continue to spread and increase protection even into the subsequent birthing season, and is less sensitive to timing than the transferable vaccine. Overall, we find that although the duration of self-dissemination influences the effectiveness of self-disseminating vaccines, it has little impact on the optimal timing of vaccine introduction: it is generally best to distribute the transmissible vaccine during the birthing season and the transferable vaccine slightly after the peak of the birthing season. Similarly, the duration of the infectious period for the pathogen has little affect on the optimal timing of vaccine delivery, although longer infectious periods decrease the vaccines ability to reduce pathogen prevalence (Supplemental Figure 3).

**Figure 4:**
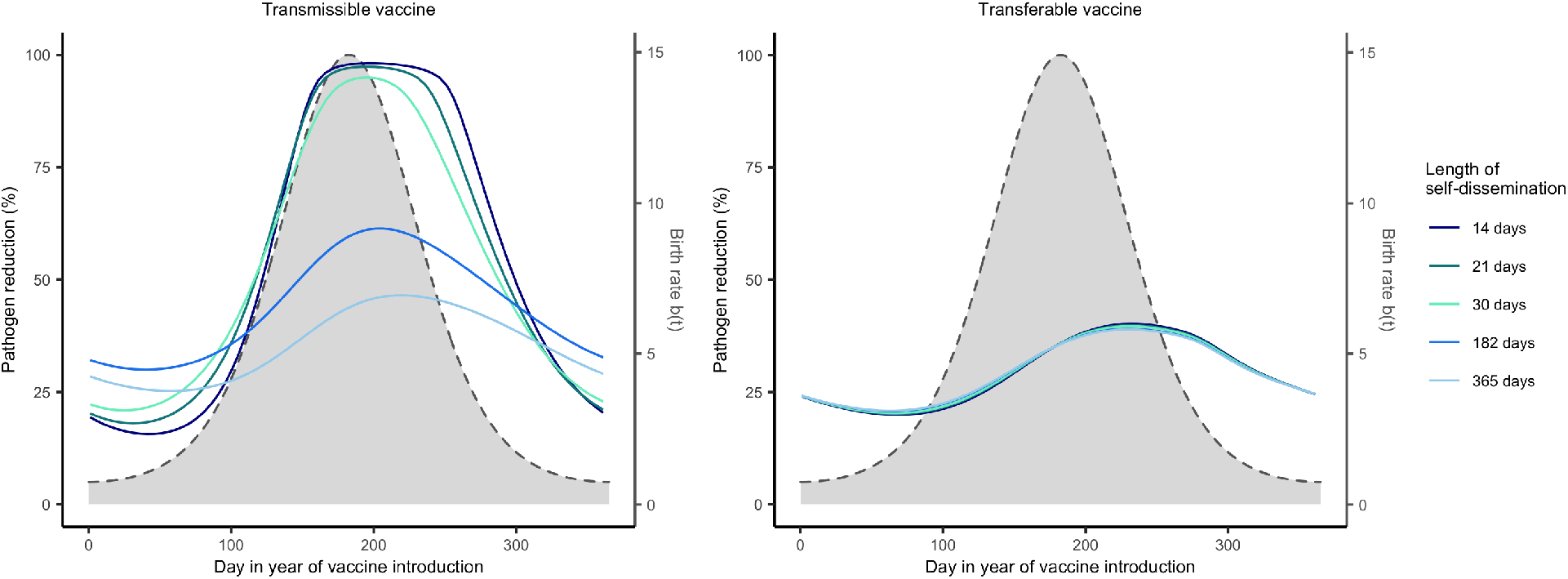
Level of pathogen reduction achieved across various times of vaccination with different vaccine recovery rates indicated by the different colors. The vaccine recovery rate controls the length of time that the vaccine can disseminate to other individuals in the population. Solid lines represent the level of pathogen reduction achieved for a given date of vaccine introduction. The grey region outlined by the dashed lined represents the seasonal birthing season where day 1 corresponds to the first day of the birthing season. The remaining parameters are: an average population size of 2000 individuals 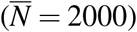, *s* = 3, an average lifespan of 1 year (*d* = 1/365), *R*_0,*V*_ = 1.5, *R*_0,*P*_ = 2, 250 vaccines are distributed each year (*N_V_* = 250), individuals remain infectious with the pathogen for 21 days on average (*γ_P_* = 21^−1^), the transferable vaccine is groomed off individuals after 6 days on average (*α* = 1/15000, and the pathogen is non-virulent (*v* = 0).

### 4.4 Reservoir life history modulates the importance of vaccine timing

We investigated how reservoir life history influences the importance of vaccine timing by adjusting average lifespan and the seasonality of reproduction. Our results demonstrate that the importance of vaccine timing decreases as average lifespan increases and has little impact when average lifespan exceeds 3 years (Figure 5). This occurs because long-lived reservoir species have a reduced rate of population turnover such that immune individuals persist within the population rather than being replaced by large quantities of susceptible individuals during the seasonal birth pulse. Even among hosts with highly synchronous births, but long lifespans, timing the delivery of vaccine made little difference in the level of pathogen reduction achieved due to long lived hosts having overall lower birth rates (Supplemental Figure 5). For those hosts with relatively brief lifespans (e.g., < 3 years), seasonality increases the importance of timing and the effectiveness of the vaccination campaign (Figure 6). This occurs because reproductive seasonality concentrates births and creates periods of time where large numbers of susceptible individuals circulate within the reservoir population. This creates opportunities for a self-disseminating vaccine to spread to a large number of individuals if its introduction is well-timed. This effect is magnified for transmissible vaccines because of their increased potential for self-dissemination.

**Figure 5:**
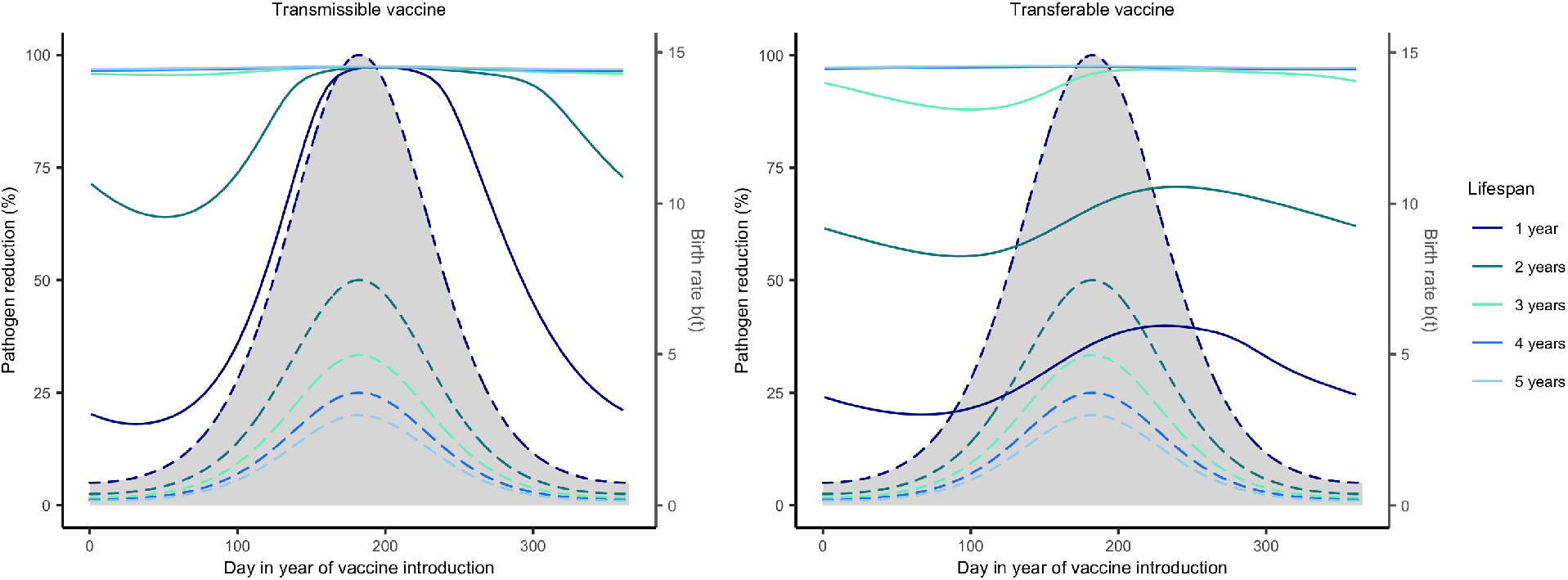
Level of pathogen reduction achieved across various times of vaccination with different average host lifespans indicated by the different colors. Each color corresponds to the host lifespan as indicated by the legend. Solid lines represent the level of pathogen reduction achieved for a given date of vaccine introduction. Dashed lines and the grey region beneath them represent the seasonal birthing season. Day 1 corresponds to the first day of the birthing season as well as the first possible day of vaccine introduction. The remaining parameters are: an average population size of 2000 individuals 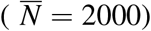, *s* = 3, *R*_0,*V*_ = 1.5, *R*_0,*P*_ = 2, 250 vaccines are distributed each year (*N_V_* = 250), individuals can disseminate vaccine for 21 days on average (*γ_V_* and *γ_g_* = 21^−1^), individuals remain infectious with the pathogen for 21 days on average (*γ_P_* = 21^−1^), the transferable vaccine is groomed off individuals after 6 days on average (*α* = 1/15000, and the pathogen is non-virulent (*v* = 0).

**Figure 6:**
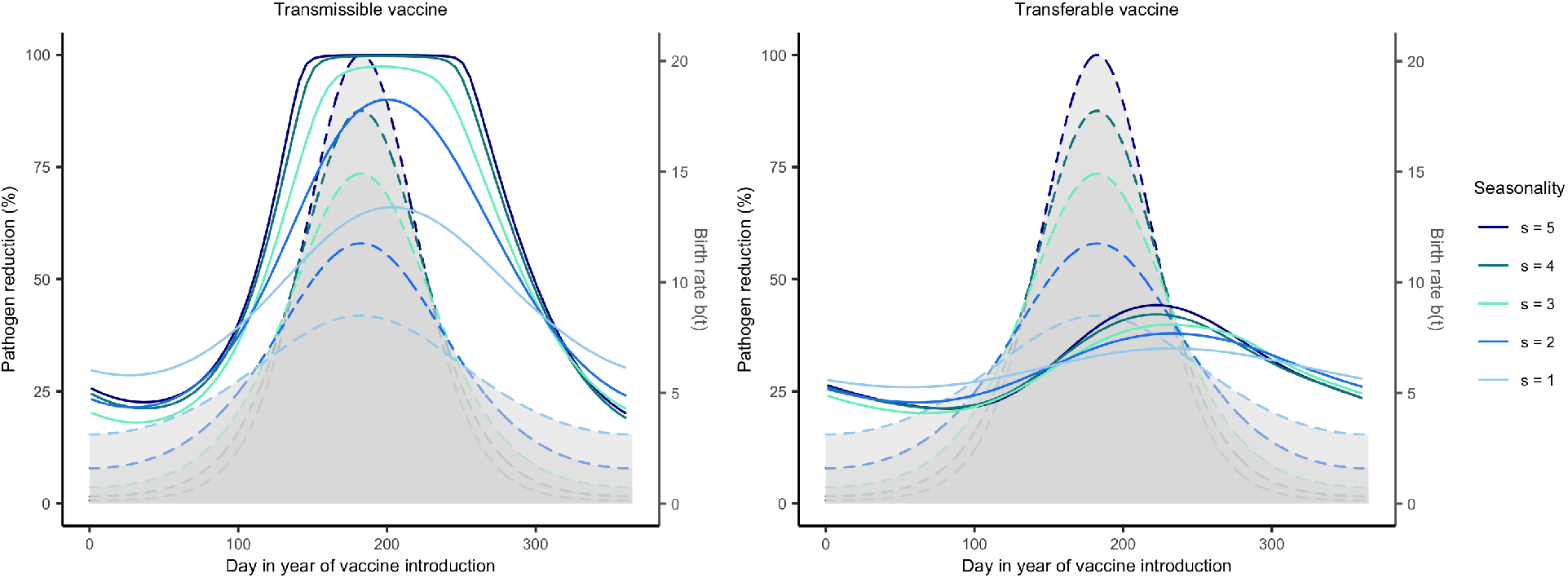
Level of pathogen reduction achieved across various times of vaccination for varying levels of synchronous births. Low s or low synchrony implies births occur over a large amount of time whereas high s or high synchrony implies all births occur over a very short time frame. Solid lines represent the level of pathogen reduction achieved for a given date of vaccine introduction. The grey region outlined by the dashed colored lines represent the seasonal birthing season for the respective parameter regime shared with the solid lines. Day 1 corresponds to the first day of the birthing season as well as the first possible day of vaccine introduction. The remaining parameters are: an average population size of 2000 individuals 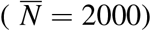, an average lifespan of 1 year (*d* = 1/365), *R*_0,*V*_ = 1.5, *R*_0,*P*_ = 2, 250 vaccines are distributed each year (*N_V_* = 250), individuals can disseminate vaccine for 21 days on average (*γ_V_* and *γ_g_* = 21^−1^), individuals remain infectious with the pathogen for 21 days on average (*γ_P_* = 21^−1^), the transferable vaccine is groomed off individuals after 6 days on average (*α* = 1 /15000, and the pathogen is non-virulent (*v* = 0).

## 5 Case study results

### 5.1 *Mastomys natalensis* – Lassa virus

We studied simulated vaccination campaigns of both the transmissible and transferable vaccine targeting Lassa virus in *M. natalensis* using the parameters described in the methods section. These simulations demonstrate that Lassa virus prevalence within the reservoir population is maximally reduced when vaccines are introduced shortly after the peak of the birthing season (Figure 7). For a transmissible vaccine with an *R*_0,*V*_ = 1, this translates into a reduction in LASV prevalence of 57% if the vaccine is introduced at the optimal time but only 37.5% if introduced before the birthing season and a transferable vaccine with *R*_0,*V*_ = 1 could achieve a 52% reduction in pathogen prevalence if timed correctly in contrast to a 25% reduction in pathogen prevalence if delivered too early. These results assume a recombinant vector transmissible vaccine created from a herpesvirus vector that causes long-term chronic infections. A transmissible vaccine constructed from a vector that generates short-term acute infections would be even more sensitive to accurate timing.

**Figure 7:**
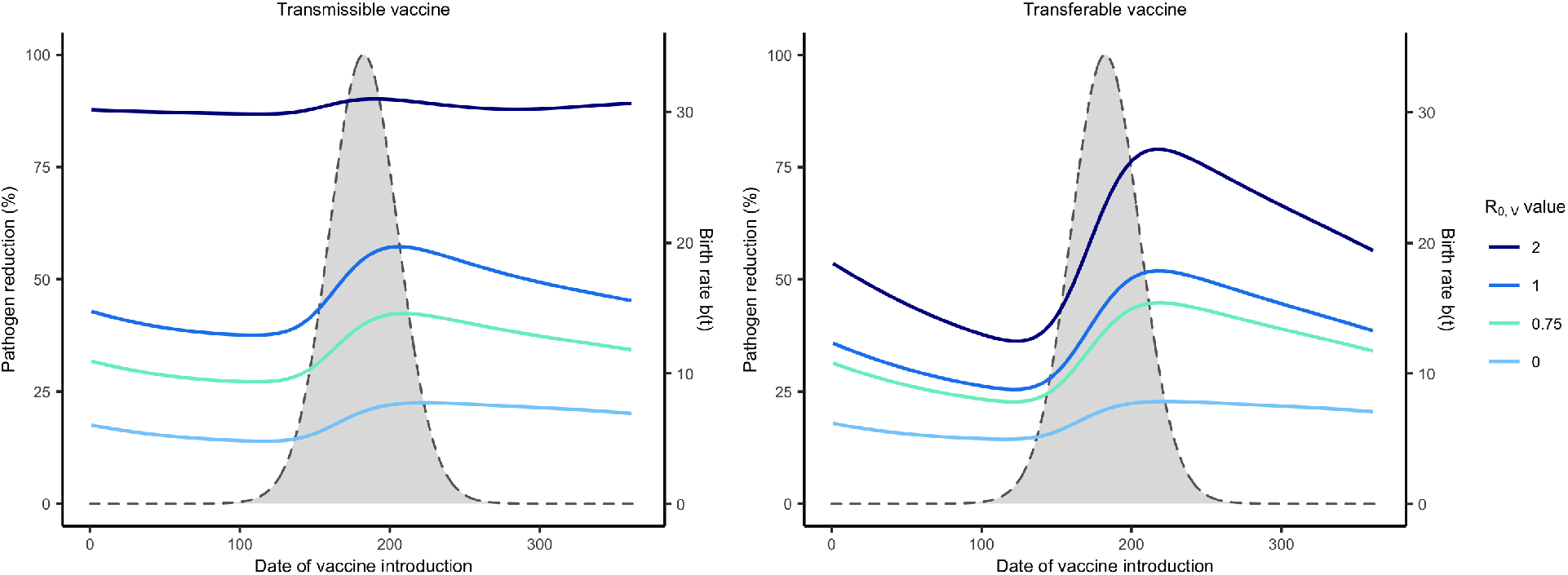
Specific example for *M. natalensis* that describes the level of pathogen reduction achieved across various times of vaccination with different vaccine *R*_0_ values indicated by the different colors. Solid lines represent the level of pathogen reduction achieved for a given date of vaccine introduction. The grey region outlined by the dashed line represents the seasonal birthing season where day 1 corresponds to the first day of the birthing season. The remaining parameters used were: an average population size of 2000 individuals 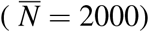, *s* = 13.078, an average lifespan of 1 year (*d* = 1/365), *R*_0,*P*_ = 1.5, 200 vaccines are distributed each year (*N_V_* = 200), individuals can disseminate the transferable vaccine for 2 days on average (*γ_g_* = 2^−1^), individuals remain infectious with the transmissible vaccine for their entire life (*γ_V_* = 0), individuals remain infectious with the pathogen for 21 days on average (*γ_P_* = 21^−1^), the transferable vaccine is groomed off individuals after 6 days on average (*α* = 1/15000, and the pathogen is non-virulent (*v* = 0).

### 5.2 *Desmodus rotundus* – Rabies virus

In addition to *M. natalensis*, we studied simulated vaccination campaigns using both transmissible and transferable vaccines targeting Rabies virus in *D. rotundus*. Simulations used the parameters described in the methods section. Our simulations demonstrate that both types of self-disseminating vaccines could substantially reduce viral prevalence within the bat population regardless of when they are distributed relative to the birthing season (Figure 8). Specifically, a transmissible vaccine with an *R*_0,*V*_ = 1 can achieve 93% reduction in rabies virus prevalence and a transferable vaccine with *R*_0,*V*_ = 1 could achieve a 96% reduction in pathogen prevalence. The transferable vaccine achieves a higher level of pathogen reduction here due to the shorter duration of self-dissemination, where as the transmissible vaccine causes lifelong infection of the vaccine. As seen previously in Figure 4, longer durations of self-dissemination lead to lower levels of pathogen reduction because – holding *R*_0,*V*_ constant – the vaccine must have a lower transmission rate. The large reductions in pathogen prevalence and the insensitivity to timing of vaccine delivery seen here for both types of vaccine are due to the substantially longer lifespan of *D. rotundus* compared to *M. natalensis*. As discussed above, organisms with longer lifespans are less sensitive to timing because these populations have low influxes of susceptible individuals each year. In contrast, short-lived organisms have high influxes of susceptible individuals which lead to a large number of individuals in the population being susceptible to the pathogen. In addition to *Desmodus rotundus* having a longer lifespan, rabies virus infection in bats can be fatal, and this may be another reason for the increased level of pathogen reduction seen here in contrast to the rodent population with Lassa virus. Specifically, we found that increasing levels of virulence can increase the level of pathogen reduction that can be achieved, and suspect that to be because pathogen mortality leads to a decrease in the number of individuals in the population that vaccines may be wasted on, see Supplemental Figure 4 for more details.

**Figure 8:**
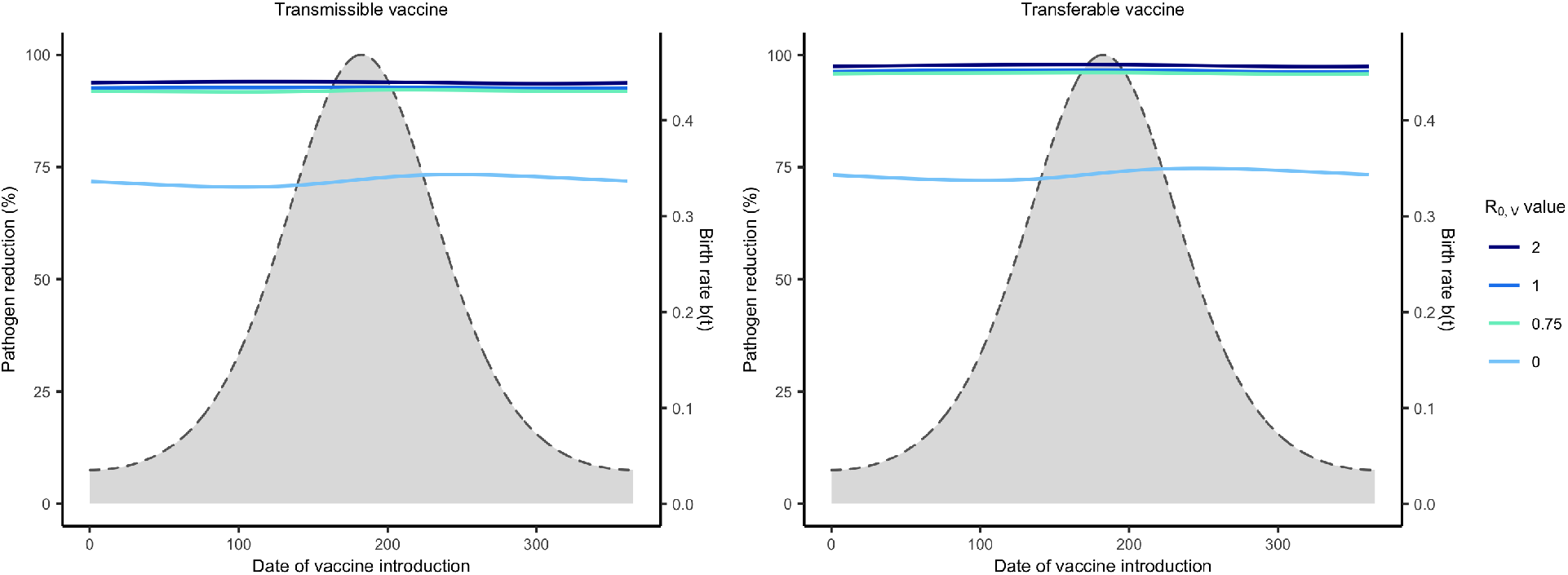
Specific example for *D. rotundus* on the level of pathogen reduction achieved across various times of vaccination with different vaccine *R*_0_s indicated by the different colors. Solid lines represent the level of pathogen reduction achieved for a given date of vaccine introduction. The grey region outlined by the dashed lined represents the seasonal birthing season where day 1 corresponds to the first day of the birthing season. The remaining parameters used were: an average population size of 240 individuals 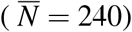, *s* = 2.59, an average lifespan of 3.5 years (*d* = 1/(365 × 3.5), *R*_0,*P*_ = 1.5, 24 vaccines are distributed each year (*N_V_* = 24), individuals can disseminate the transferable vaccine for 7 days on average (*γ_g_* = 2^−1^), individuals remain infectious with the transmissible vaccine for their entire life (*γ_V_* = 0), individuals remain infectious with the pathogen for 21 days on average (*γ_P_* = 21^−1^), the transferable vaccine is groomed off individuals after 6 days on average (*α* = 1/15000, and the pathogen is virulent (*v* = 0.005).

## 6 Discussion

We have used mathematical models of self-disseminating vaccines to evaluate how the timing and duration of vaccine distribution influences the impact of vaccination campaigns targeting seasonally fluctuating wildlife populations. Our results demonstrate that self-disseminating vaccines increase protection relative to traditional vaccines but that the magnitude of this increase can be sensitive to the timing of vaccine distribution. This is particularly true for transmissible vaccines that transmit only weakly and for transferable vaccines. Sensitivity to timing is also most important for reservoir species with short lifespans and distinct birthing seasons. In these scenarios, it is generally best to distribute vaccine shortly after the peak in the reservoir birthing season. This general result mirrors previous findings for traditional, non self-disseminating wildlife vaccines from Schreiner et al. (2020), but clarifies how the magnitude of the effect depends on the type of self-disseminating vaccine and its specific properties.

An important result that emerges from our work is that transferable vaccines are more sensitive to timing than are transmissible vaccines. This occurs primarily because transmissible vaccines can generate self-sustaining chains of transmission whereas transferable vaccines cannot. Thus, transferable vaccines can spread only to susceptible individuals at the time of vaccine introduction. In contrast, transmissible vaccines can be introduced earlier and yet still reach individuals that will be born later through persistent chains of vaccine transmission. This insensitivity to timing is greatest for highly contagious transmissible vaccines that generate long chains of transmission.

The importance of our results for real world applications depends on reservoir lifespan and the extent to which reservoir reproduction is seasonal. As demonstrated by our general and case study results, the lifespan of hosts has a large effect on the sensitivity to seasonality because it influences population turnover. For example, our results show that the success of attempts to vaccinate *M. natalensis*, the reservoir of Lassa virus, may be very sensitive to timing because the reservoir has a short lifespan. This sensitivity arises because rapid turnover within the reservoir population leads to a large, seasonal influx of susceptible individuals. In contrast, our results show that efforts to vaccinate the vampire bat, *D. rotundus*, are not particularly sensitive to timing due to the long lifespan of the reservoir. In long-lived populations like these, population turnover is low and the seasonal influx of newly born susceptible individuals relatively small. Although we have illustrated the relevance of our general results using the specific examples of Lassa virus and rabies virus, these general results have broad implications for efforts to vaccinate reservoir animals against other important human pathogens. For instance, hantaviruses, such as Sin Nombre virus, also have reservoir species that are short-lived and have seasonal reproduction (Mills et al., 1999). In these cases, our results suggest that vaccination efforts will need to be well-timed and carefully planned to achieve maximum effectiveness.

Although we believe our results are broadly generalizable, they do rest on three important assumptions. First, we have ignored reservoir age structure. Age structure may influence the number of actively foraging animals in the population, leading to different rates of vaccine uptake in young versus adult individuals, as has been seen for raccoon oral rabies vaccination campaigns (Mainguy et al., 2012). Optimal timing may change from what we predict in such a scenario because we assume newborn susceptible individuals consume vaccine. Second, we do not take maternal antibodies into consideration. Presence of maternal antibodies has been demonstrated in foxes, rodents, and bats and may prevent juveniles from developing a robust and long-lasting immune response to the vaccine (Müller et al., 2001; Mariën et al., 2019; Constantine et al., 1968; Shankar et al., 2004). This may lead to wasted vaccine if vaccines are distributed while maternal antibodies interfere with vaccine effectiveness (Zhi Q. and Hildegund C.J., 1992). For instance, antibodies in red foxes have been shown to persist for 8 weeks (Müller et al., 2001). In such cases, vaccination may need to be delayed relative to what we predict here to avoid interference between vaccine and antibodies. In general, the need to avoid interference with maternal antibodies may narrow the window of opportunity for effective vaccine distribution and make timing even more important than our results suggest.

Self-disseminating vaccines make vaccinating hard-to-reach wildlife populations more feasible. Our results show that optimizing the timing and duration of vaccine delivery can make or break the success of a vaccination program in fluctuating wildlife populations with high levels of population turnover. These results further demonstrate the importance of understanding the population ecology of wildlife species prior to implementing vaccination campaigns using self-disseminating vaccines.

## 7 Acknowledgements

We thank Jim Bull for helpful comments regarding this work. This work was supported by NSF DEB 1450653 (SLN and CHR), NIH R01GM122079 (SLN and CHR), and DARPA D18AC00028 (SLN).

## 8 Appendix

### 8.1 Setting the birth scaling constant *k*

In our simulations, the scaling constant *k* in the birthing function is determined by the user-specified values *d*, *s*, and 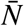. To solve for the value of *k*, we first rewrite the birthing function as

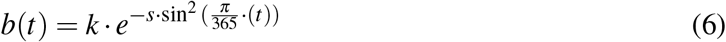

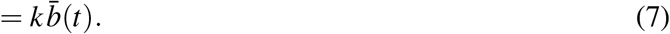

The differential equation that describes the host population size in the absence of any infectious agent is

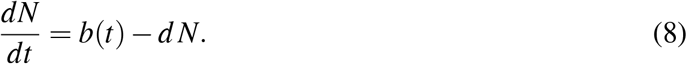

Let *N*\*(*t*) denote the *T*-periodic solution of Eq (8) with mean value 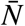. Then

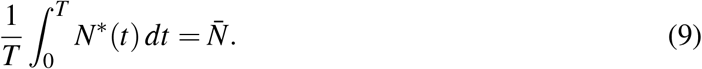

This implies

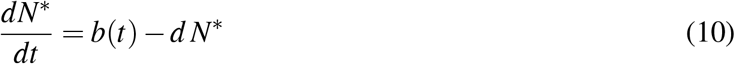

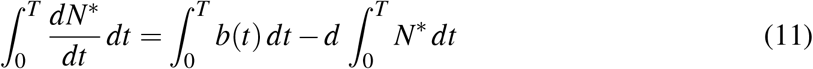

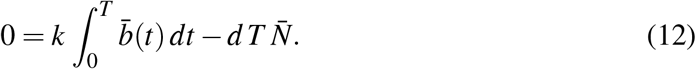

The left hand side of Eq (12) is zero because *N\** is T-periodic. Thus, we have

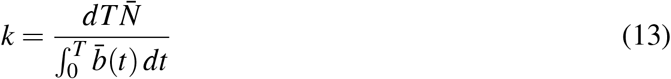

Thus, for a specified *d*, *s*, *T*, and 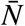, Eq (13) can be numerically integrated to solve for the implied value of *k*.

### 8.2 Derivation of *R*_0_

In this section, we derive an expression for the basic reproduction number, notated *R*_0_, that describes the average number of new infections that result when a single infected individual is introduced at a random time into a stably cycling population of susceptible hosts. We keep our derivation broad so as to simultaneously derive the relevant *R*_0_ for the pathogen, transmissible vaccine, and transferable vaccine, under both density and frequency-dependent transmission.

Let *N*\*(*t*) denote the T-periodic limit cycle that describes a population of susceptible hosts in the absence of infection and vaccination. We assume that *N*\*(*t*) >> 1 so that the susceptible population is not significantly depleted by the infection process. Let *β*, *γ, v* denote the transmission rate, recovery rate, and the virulence rate of the infectious agent. Let *C*(*N*) describe how the percapita rates of host interaction scale with population size: under a density-dependent scenario, *C*(*N*) = *N*, while under a frequency-dependent scenario, *C*(*N*) = 1 (Keeling and Rohani, 2011). For the transferable vaccine, we assume that grooming interactions scale with population size in the same way as infectious contacts. Thus, *αC*(*N*) describes the rate at which vaccine is groomed off gelled individuals in a population of size *N*.

When a single infected host is introduced into a susceptible population described by *N*\*(*t*), the rate of new infections at time *t* is *βC*(*N*\*). Here, we omit the dependence of *N\** on *t* to simplify notation. Depending on the infectious agent being described, this infection rate continues until the initial infected host dies due to natural mortality (at rate *d*), dies due to pathogen virulence (at rate *v*), recovers from infection (at rate *γ*), or in the transferable vaccine case, leaves the infectious class due to grooming of gel at rate *αC*(*N*\*). Note that *α* = 0 in the case of the pathogen or transmissible vaccine.

Let *t*_0_ denote the time at which the infected individual is introduced. The total number of new infections caused by the infected individual is obtained by integrating the infection rate (*βC*(*N*\*)) multiplied by the probability that the individual has not recovered or died from time *t* = *t*_0_ to time *t* = ∞.

To find the probability that the individual has not lost infectiousness status, let *P*(*t, t*_0_) denote the probability that the individual is still infectious at time *t* > *t*_0_. We assume that *P*(*t, t*_0_) is described by a Poisson process with probabilistic rates at which infectiousness is lost due to natural death (*d*), degradation of vaccine or recovery (*γ*), mortality due to pathogen virulence (*v*), or grooming (*αC*(*N*\*)). Then for time *t* > *t*_0_, and a small time interval Δ*t*, *P*(*t, t*_0_) satisfies

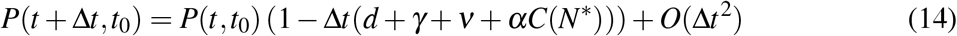

with initial condition *P*(*t*_0_, *t*_0_) = 1. Here, *O*(Δ*t*^2^) denotes terms in Eq (14) that become negligible in the limit as Δ*t* approaches zero. In words, Eq (14) describes how the probability of the individual still being infectious at time *t* + Δ*t* is approximately equal to the probability that the individual was infectious at time *t*, multiplied by the probability that the individual’s infectious status has not changed in the interval (*t, t* + Δ*t*).

By rearranging terms in Eq (14) and taking the limit as Δ*t* approaches zero, we derive the continuous time differential equation

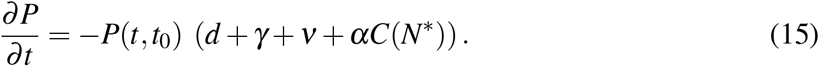

Dividing both sides of Eq (15) by *P*(*t, t*_0_) and integrating over *t* from time *t*_0_ yields the probability that an initial infected individual introduced at time *t*_0_ is still capable of infecting others at time *t*:

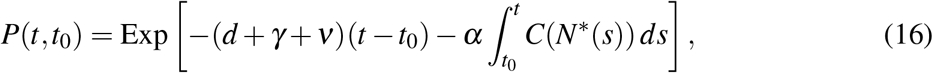

where Exp[x]= *e^x^* denotes the exponential function.

With Eq (16) in hand, we can express the total number of new infections caused by the introduced infected individual as

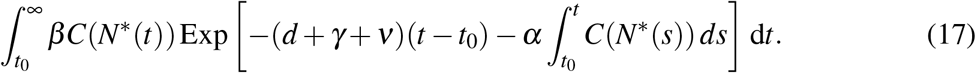

Eq (17) highlights that, because the population size *N*\*(*t*) is non-constant, the number of new infections is a function of the time *t*_0_ at which the infected individual is introduced. In order to find the average number of new infections generated by an infected that is introduced at a randomly chosen time, we integrate Eq (17) with respect to *t*_0_ over the interval [0, *T*], and divide by 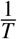. Note that because *N*\*(*t*) is T-periodic, averaging over introduction times that are outside the interval [0, *T*] is redundant. Consequently, we have

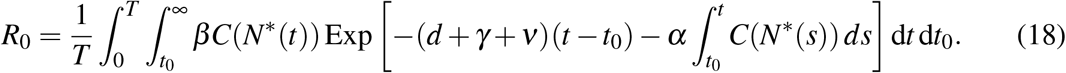

#### 8.2.1 Transferable vaccine and density-dependent scaling of host interactions

In the case of the transferable vaccine and density dependent host interactions, virulence is absent so we set *v* = 0. In addition, *α* ≠ 0 and *C*(*N*\*) = *N\** so the integral described by Eq (18) is difficult to simplify by the presence of the antiderivative of *N*\*(*t*) in the exponent.

#### 8.2.2 Transmissible vaccine and pathogen or frequency-dependent scaling of host interactions

In the case of the transmissible vaccine and the pathogen, or when interactions are frequency dependent, the expression for *R*_0_ in Eq (18) can be simplified. In all of these cases, the double integral described by Eq (18) can be simplified by using the change of coordinates *u* = *t*, *w* = *t* – *t*_0_. This change of coordinates needs to be applied to three terms in the above integral: the area differential d*t* d*t*_0_, the limits of integration, and the integrand (Stewart, 2012).

Let *X*(*u, w*) = (*u, u – w*) denote the vector-valued function that converts (*u, w*) coordinates into (*t, t*_0_) coordinates. Then the area differential d*t* d*t*_0_ is equal to |*DX*| d*u* d*w*, where *D* denotes the Jacobian operator with respect to *u* and *w*, and | · | denotes the determinant. Because |*DX*| = 1, we have d*t* d*t*_0_ = d*u* d*w*. The region of integration in the (*u, w*) plane can be found by drawing the region of integration in the (*t, t*_0_) plane, and identifying boundary lines with their analogue in the (*u, w*) plane (Figure 9). Finally, the integrand is transformed by the substitution *t* → *u* and *t – t*_0_ → *w*.

**Figure 9:**
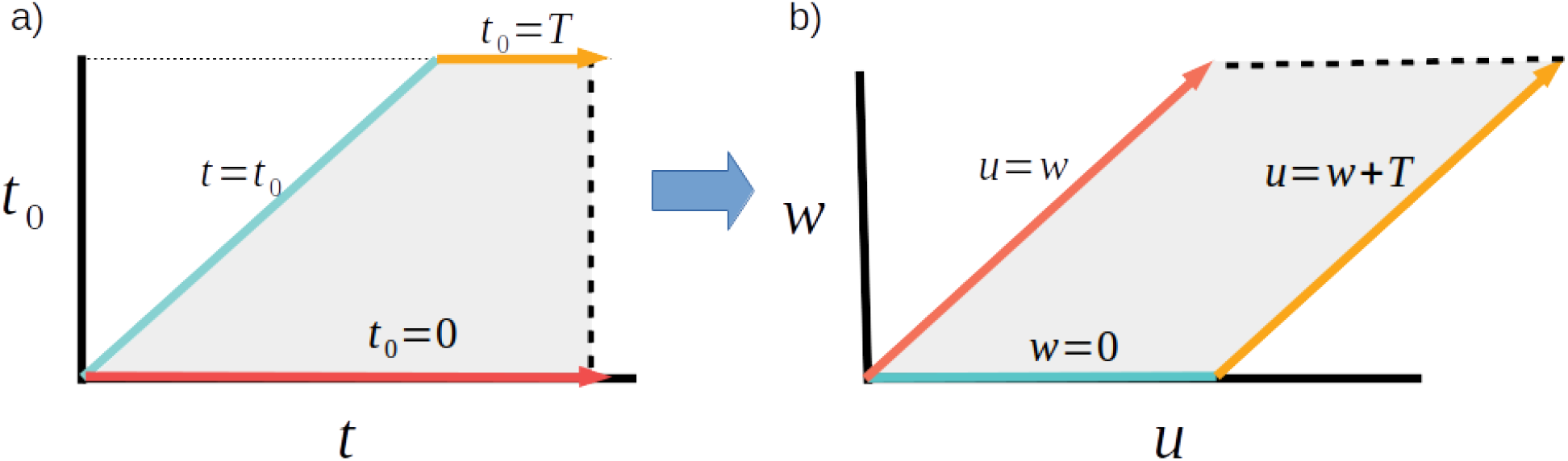
Region of integration (gray) of Eq (18) in the (*t, t*_0_) plane (a). When *α* = 0, the calculation of *R*_0_ is simplified by transforming the region into the (*u, w*) plane (b). The dashed boundary lines indicate that the region continues out to infinity. Boundary lines and their transforms are identified by the same color.

We first evaluate the case when host interactions are density-dependent (*C*(*N*) = *N*). When *α* = 0 these substitutions allow us to transform the integral in Eq. (18) and evaluate as follows:

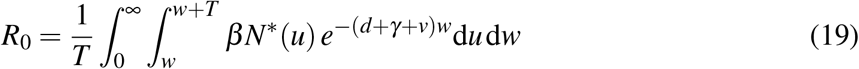

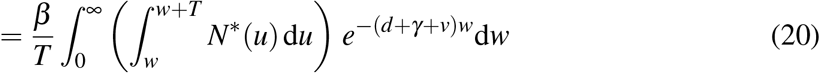

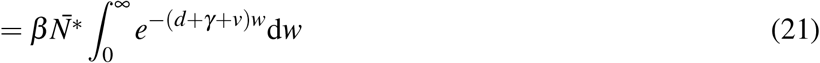

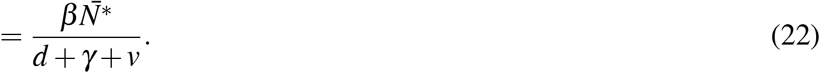

Here, 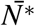 denotes the average population size over a single period *T*. Virulence *v* is possibly nonzero for the pathogen’s *R*_0_, and virulence is set to zero for the transmissible or transferable vaccines’ *R*_0_’s.

When the rate of interactions is frequency-dependent and for general *α* ≥ 0, the sequence of equations 19 - 22 can be applied in a similar manner to obtain

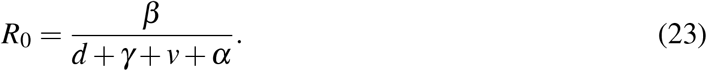

### 8.3 Setting the transmission rate *β*

Equations (18), (22), and (23) are used to define the transmission rate *β* that corresponds to specific values of *R*_0_ in our simulations. For a given simulation and infectious agent, we define an average population size 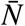, death rate *d*, virulence *v*, recovery rate *γ*, gel grooming rate *α*, and basic reproduction number *R*_0_. In the case of a density dependence and for the transmissible vaccine or pathogen, Eq (22) can then be used to solve for the value of *β* that is implied by the user-defined parameters. If the host interaction rate is frequency dependent, Eq (23) is used to derive *β*.

The density dependent, transferable vaccine case is more difficult because we need the solution of *N*\*(*t*) to evaluate Eq (18). To this end, we first solve for the value of *k* using Eq (13) and parameters specified by the user. *k*, in turn, is used to define the birthing rate *b(t*). Next, we obtain a numerical approximation of *N* (t*) by simulating the population equation Eq (8). Specifically, we simulate Eq (8) for 10 years to allow the solution to converge to the stable limit cycle *N*\*(*t*). Next, we use the function “approxfun” in R to approximate the stable limit cycle *N*\*(*t*). Finally, we use these numerical approximations to evaluate the double integral of Eq (18) and solve for the value of *β* that is implied by a user specified *R*_0_. All integration was performed in the statistical language R using the deSolve package (Soetaert et al., 2010).

### 8.4 Estimating seasonality parameter for case studies

In this section, we describe how the seasonality parameter *s* was parameterized for the case studies on *Mastomys natalensis* and *Desmodus rotundus*.

#### 8.4.1 Mastomys natalensis

We use data on trapping success of *M. natalensis* in Guinea to broadly estimate the seasonality parameter *s* (Fichet-Calvet et al., 2007). This study contains time series of trap success from two towns. Because *M. natalensis* is typically associated with human habitation, we use the within-house trap success as a relative measure of *M. natalensis* population size. We choose a value of *s* so that, when the average population size in our simulation is 2000 rodents, the ratio of the maximum and minimum population size from our model matches the ratio of the maximum to minimum trap success from these time series data. Figure 2 of the study implies that this ratio is approximately two (Fichet-Calvet et al., 2007). With this ratio in hand, we use the “optimize” function in R and the population demography model described by Eq (8) to find the value of *s* that minimizes the squared error between the simulated maximum:minimum population ratio (after a 101 year burn-in period) and the estimated true ratio of two. This method yields a value of *s* = 13.078.

#### 8.4.2 Desmodus rotundus

To parameterize the birth function for *Desmodus rotundus*, we choose a value of the seasonality parameter *s* that matches the ratio of the maximum birth rate to the minimum birth rate. We do this by using the analytical form of the maximum and minimum of the birth function. For a given year *n* (a positive integer), the function *b*(*t*) reaches its seasonal maximum *k* at *t* = 365*n* and minimum *k* · exp –*s* when 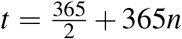. We then use the ratio of the maximum and minimum birth rate, respectively referred to as “max” and “min”, to estimate the seasonality parameter *s*:

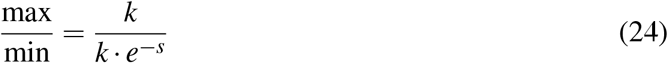

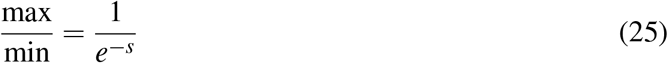

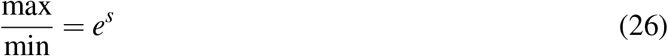

Thus, we find

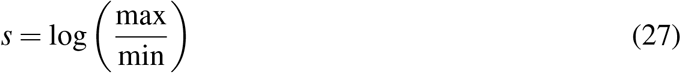

With Eq (27) in hand, we can use data on the estimates of birth rate throughout the year to estimate *s*.

Estimates for birth rates for *Desmodus rotundus* have come from data on lactating females. Specifically, studies in Argentina on vampire bats found a direct relationship between the number of lactating females and the number of births in the population (Lord et al., 1976). We use estimates and the method outlined above to find an estimate for s. Based on Figure 1 from Lord et al. (1976) and Figure S1 from Blackwood et al. (2013) we estimate that the ratio of the maximum:minimum birth rate is 40:3. With these values, Eq (27) implies *s* = 2.59.

